# PyDesigner: A Pythonic Implementation of the DESIGNER Pipeline for Diffusion Tensor and Diffusional Kurtosis Imaging

**DOI:** 10.1101/2021.10.20.465189

**Authors:** Siddhartha Dhiman, Joshua B Teves, Kathryn E Thorn, Emilie T McKinnon, Hunter G Moss, Vitria Adisetiyo, Benjamin Ades-Aron, Jelle Veraart, Jenny Chen, Els Fieremans, Andreana Benitez, Joseph A Helpern, Jens H Jensen

## Abstract

PyDesigner is an open-source and containerized Python software package, adapted from the DESIGNER pipeline, for diffusion weighted magnetic resonance imaging preprocessing and tensor estimation. PyDesigner combines tools from FSL and MRtrix3 to reduce the effects of signal noise and imaging artifacts on multiple diffusion measures that can be derived from the diffusion and kurtosis tensors. This publication describes the main features of PyDesigner and highlights its ease of use across platforms, while examining its accuracy and robustness in deriving commonly used diffusion and kurtosis metrics.

## INTRODUCTION

Diffusion MRI (dMRI) is widely applied for the noninvasive study of microstructural properties in the brain. While many dMRI methods have been proposed, two of the most commonly used are diffusion tensor imaging (DTI) and diffusional kurtosis imaging (DKI). These techniques are closely related, with DKI being a generalization of DTI that includes quantification of diffusional non-Gaussianity (Jensen and Helpern, 2010). Both provide a variety of scalar diffusion measures and enable the construction of white matter fiber tractography. An important advantage of DTI and DKI is that they have a solid foundation in diffusion physics so that their validity does not rely on detailed assumptions regarding tissue microstructure (Basser, 2002; Jensen et al., 2005). This allows DTI and DKI to be applied throughout the brain and body for both healthy and diseased subjects of any age.

Because raw diffusion weighted images (DWIs) are degraded by multiple factors, including signal noise, motion, Gibbs ringing, and eddy current distortion, preprocessing should be employed prior to calculation of any diffusion quantities (Le Bihan et al., 2006). Preprocessing of DWIs is now highly developed, and several popular software packages are freely available for performing the various preprocessing steps. However, combining these steps into a single pipeline that gives consistent results is challenging both because there are a number of user defined settings that must be adjusted depending on the details of the dMRI acquisition and because the order in which the preprocessing steps are performed affects the outcome. For this reason, the Diffusion parameter EStImation with Gibbs and NoisE Removal (DESIGNER, GitHub: NYU-DiffusionMRI/DESIGNER) pipeline was proposed in order to optimize, standardize, and streamline the preprocessing for DWIs. DESIGNER relies on FSL, MRtrix3, MATLAB, and Python to create a seamless and complete DWI processes – one that encompasses image correction through preprocessing and diffusion/kurtosis tensor estimation (Ades-Aron et al., 2018). With control flags to toggle preprocessing steps on or off, DWI corrections can be performed selectively. DESIGNER always preprocesses in a specific manner – (i) Marchenko-Pastur principal component analysis (MP-PCA) denoising, (ii) Gibbs ringing correction, (iii) echo-planar imaging (EPI) distortion correction, eddy current correction, motion correction, and outlier replacement, (iv) B1 bias field correction, (v) brain mask generation, (vi) smoothing, (vii) Rician noise bias correction, and (viii) b0 normalization. Preprocessing in this specific order improves both accuracy and the effective signal-to-noise ratio (SNR) (Ades-Aron et al., 2018).

Implementing DESIGNER across platforms is challenging because of differences in operating systems and environment settings. In particular, the fact that DESIGNER is mainly written in MATLAB creates significant portability issues arising from complicated configuration requirements needed to enable Python-MATLAB interfacing. Moreover, reproducibility of outputs can be compromised from different combinations of MATLAB, Python and dependency versions. For this reason, we have developed PyDesigner, which is entirely Python based. Not only does this allow for seamless preprocessing, but it also allows PyDesigner to be incorporated into a Docker container that greatly enhances portability and reproducibility. Moreover, by replacing the MATLAB code, PyDesigner avoids all licensing fees and improves accessibility.

The purpose of this paper is to describe the main features of PyDesigner and its implementation. PyDesigner augments the hands-free approach introduced by DESIGNER, adds several new features, and incorporates tools from FSL and MRtrix3 to perform preprocessing. Standard mathematical Python libraries such as Numpy (Harris et al., 2020), SciPy (SciPy 1.0 Contributors et al., 2020), and CVXPY (Agrawal et al., 2018; Diamond and Boyd, 2016) were used to replace the MATLAB portions of DESIGNER with Python code. All PyDesigner software is open source and available at: https://github.com/m-ama/PyDesigner.

While alternative software such as Diffusion Kurtosis Estimator (DKE, Tabesh et al., 2011) and Diffusion Imaging in Python (DIPY, Henriques et al., 2021) are also available, they do not combine image correction and tensor fitting into a single-command pipeline. The DIPY package is a Python-based image correction and tensor fitting tool, whereas DKE is largely a tensor fitting tool that runs in MATLAB. Here, the core calculation of tensor fitting and associated diffusion parameter estimation of PyDesigner is compared with that of DESIGNER, DKE, and DIPY to examine computational differences and illustrate the relative performance of these four DKI analysis programs. In making this comparison, the PyDesigner preprocessing is applied in all four cases so that any differences are entirely attributable to the tensor fitting step.

## METHODS

### Workstation

All processing was performed on a custom-built workstation, equipped with 8-cores AMD Ryzen 2700x, 16 GB system memory, and Nvidia GTX 1080 running on CUDA v10.1.

### OS Information

Ubuntu 20.04 (Focal Fossa) was used with the software packages FSL v6.0, MRtrix3 v3.0.1-24-g62bb3c69, and Conda 4.8.3 with a custom Python 3.6 environment containing PyDesigner v1.0-RC8 and all dependencies.

### DWI Acquisition

An acquisition from a cognitively healthy subject in their 20s was acquired using the Siemens Prisma^fit^ 3T scanner (Siemens Healthineers, Erlangen, Germany). DTI and DKI sequences were acquired with 3 *b*-values (*b* = 0, 1000, 2000 s/mm^2^) for 10 images acquired at *b* = 0 (b0) and 30 isotopically distributed diffusion encoding directions for *b* = 1000 and 2000 s/mm^2^. This protocol was performed using single-shot, twice refocused echo-planar sequence at 3 mm isotropic resolution with echo time (TE)/repetition time (TR) = 95/4800 ms, 74×74 acquisition matrix, 42 axial slices, bandwidth of 1648 Hz/px, slice acceleration factor = 2, parallel imaging factor = 2, and anterior (A)≫posterior (P) phase encoding direction. A separate *b*0 volume in P≫A phase encoding direction was acquired for distortion correction using TOPUP (Andersson et al., 2003). All acquisitions were acquired with full Fourier coverage.

### Staging

Acquired images were converted from DICOM to NifTi-standard with *dcm2niix* v1.0.20181125 (Li et al., 2016) to generate 4D NifTi image volumes (*.nii*), gradient (*.bvec*) files, *b*-value (*.bval*)files, and accompanying BIDS sidecars (*.json).* PyDesigner seeks JSON tags *PartialFourier, PhaseEncodingSteps, AcquisitonMatrixPE* and *EchoTime* to automatically determine ideal image correction steps.

### Preprocessing

Staged files were processed with PyDesigner using the flags --*standard* for standard preprocessing; and --*rpe_pairs 1* since a single pair of reverse phase-encoded b0s were acquired for EPI correction. The full command parsed was

1. pydesigner
2. --standard \
3. --rpe_pairs 1 \
4. --verbose \
5. -o [PATH TO OUTPUT DIRECTORY] \
6. $DKI_PROTOCOL.nii,$FBI_PROTOCOL.nii

These flags yield the following preprocessing steps, in order of appearance: (1) MP-PCA denoising, (2) Gibbs ringing correction, (3) EPI distortion correction using one pair of reverse phase-encoded b0s, (4) eddy current, motion and outlier correction, (5) brain masking, (6) smoothing at 1.25 × FWHM, (7) Rician noise bias correction, (8) mean b0 volume extraction, (9) iterative reweighted linear least squares (IRLLS) outlier rejection, (10) brute-forced tensor correction, (11) constrained tensor fitting, (12) DTI and DKI scalar map extraction, in order of appearence. A visual representation of these preprocessing steps can be found in Figure 1. These steps can be grouped into image correction and tensor fitting, where the former aims to minimize noise and correct artifacts, and the latter performs computations to derive useful dMRI metrics.

**Figure 1:**
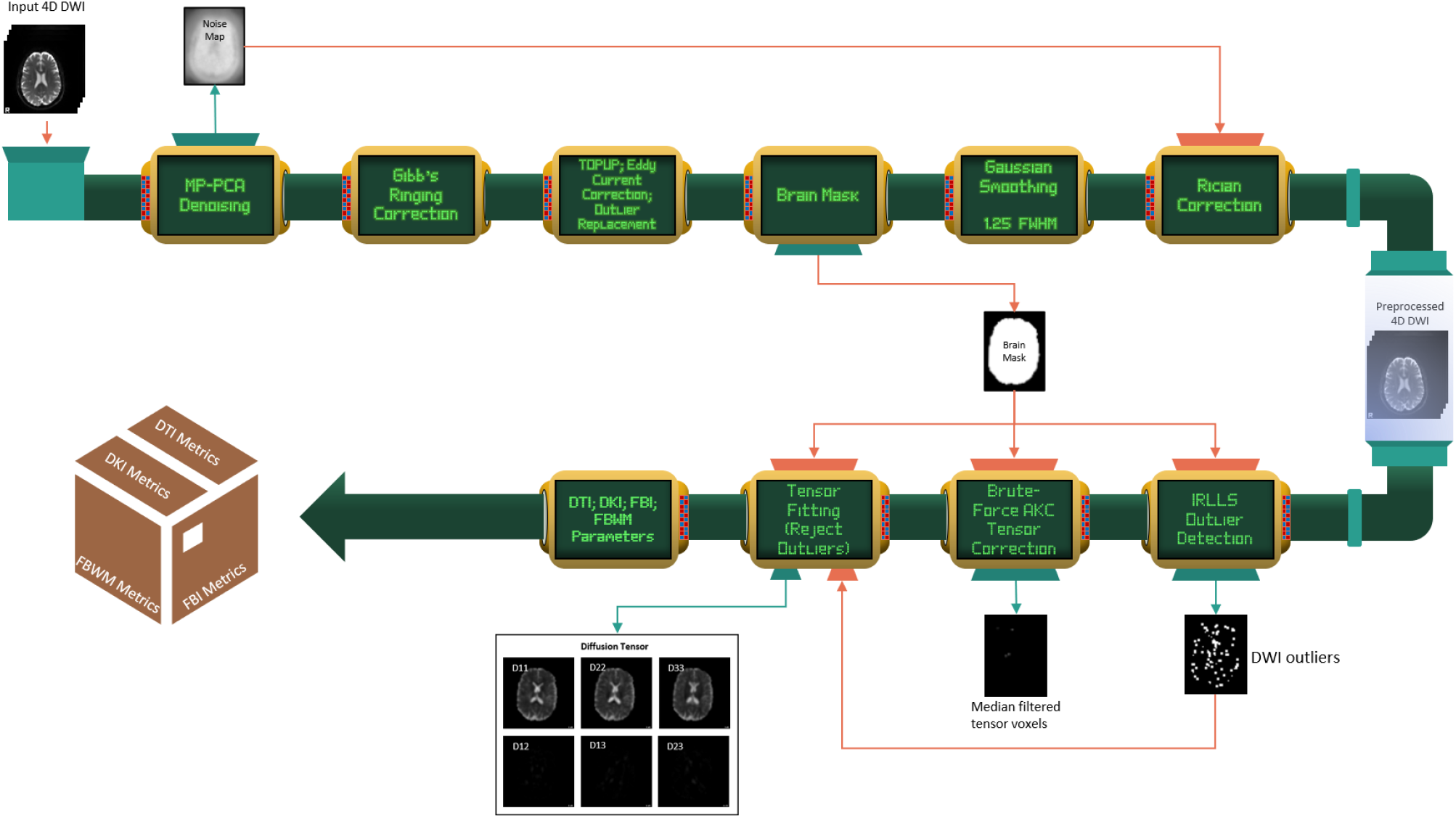
Visual representation of the PyDesigner pipeline; order of processing is clockwise. Preprocessing begins by providing an input 4D diffusion weighted image (DWI) to PyDesigner (top left), which then undergoes MP-PCA denoising to yield a noise-free 4D DWI and a 3D noise map. The noise-free 4D DWI then undergoes Gibbs ringing correction, TOPUP, eddy current correction and outlier correction. A brain mask is then computed for subsequent steps to speed up computations by performing them within the brain mask only.

#### Image Correction

##### ● MP-PCA Denoising

Preprocessing is initiated with MP-PCA denoising, using the MRtrix function *dwidenoise*, to retain noncorrelation between spatial and successive volume voxels (Veraart et al., 2016a, 2016b). Preprocessing is initiated with denoising, as the following steps introduce interpolation or reduced entropy, which can skew the underlying Marchenko-Pastur distribution and inhibit accurate noise estimation.

##### ● Gibbs ringing correction

Next, Gibbs ringing correction is introduced using the MRtrix function *mrdegibbs*, which resamples the DWI at zero-crossings of the sinc-function to remove ringing artifacts with minimal smoothing (Kellner et al., 2016). Readers should note that this correction is only applied if a DWI is acquired with full k-space coverage (full Fourier) so that sub-voxel shifts can be accurated predicted. PyDesigner automatically detemines the k-space coverage of an image using image metadata and will disable this correction if partial coverage is found. Of interest for future updates to PyDesigner, recently, a Removal of Partial-fourier Gibbs (RPG) ringing artifact method has been proposed by (Lee et al., 2021) extending the original sub-voxel shift method.

##### ● Susceptibility-induced and Eddy current correction

After the two preceding low-pass filters, susceptibility-induced distortion is corrected using a single pair of reverse phase-encoded b0s to allow rapid EPI correction without the risk of overestimating the distortion field. This is followed by motion, b-matrix rotation, eddy current and outlier correction, which results in a co-registered DWI free of outlier voxels (Andersson et al., 2016; Andersson and Sotiropoulos, 2016). This correction is applied through MRtrix’s *dwifslpreproc* function, which is a wrapper for FSL’s *topup* and *eddy* functions. Phase encoding information in the image metadata is read by PyDesigner to automate this step, so users are not required to manually specify phase encodings.

##### ● Brain mask

With all DWI volumes co-registered, a mean b0 volume is used to create a brain mask using FSL’s *bet* at 0.25 threshold for subsequent steps. Users can adjust *bet* threshold with the --*maskthr* flag or supply their own brain mask with the --*user_mask* flag.

##### ● Smoothing

Smoothing with a Gaussian kernel 1.25 × full-width half maximum (FWHM) of the voxel size is then applied to attenuate residual noise or artifacts that may have remained after prior corrections. While not entirely necessary because of smoothing-free algorithms used in previous preprocessing steps, it diminishes effects of outlier voxels. While PyDesigner defaults to 1.25 × FWHM, users are able to adjust the size of the Gaussian kernel with the --fwhm flag.

##### ● Rician noise bias correction

The final image correction is another low-pass filter to dampen the Rician noise bias generated by taking the magnitude of the raw DWIs during scanner reconstruction. The noise map derived from MP-PCA denoising is used to estimate the unbiased noise standard deviation, thus enabling an estimation of the true signal voxel intensity (Koay and Basser, 2006).

#### Tensor Fitting

##### ● IRLLS outlier detection

Diffusion tensor fitting is initiated with IRLLS to undermine skewness of data distribution, so voxels demonstrating hypo- and hyperintensities can be marked as outliers (Collier et al., 2015).

##### ● Outlier-excluded constrained tensor fitting

Voxels unmarked by IRLLS undergo a constrained and log-linearized (Veraart et al., 2013) diffusion and kurtosis tensor fitting through a quadratic program (QP), where positive apparent kurtosis (*K_app_* > 0) is defined as the default constraint (Tabesh et al., 2011). There are a total of three constraints that can be toggled on or off with the -- *fit*_*constraints* flag to limit tensor fitting such that *D_app_* > 0, *K_app_* > 0, or 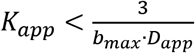 where *D_app_*is apparent diffusion coefficient, *K_app_* is apparent kurtosis coefficient, and *b_max_* is the maximum *b*-value of the data.

##### ● Brute-forced apparent kurtosis coefficient (AKC) correction

Fitted tensors undergo additional refinement by brute-forcing them across 100,000 pre-defined gradient directions to compute AKC values, where tensor voxels with AKC less than –2 or more than 10 are median filtered. Users are cautioned that this method is not yet validated and can introduce outliers in some instances. While future updates to PyDesigner are expected to deprecate AKC correction, current users may disable it with the --*noakc* flag.

##### ● Parameter extraction

Culmination of IRLLS, constrained tensor fitting, and brute-forced AKC correction yield biologically plausible tensors suitable for microstructural evaluation via DTI, DKI, fiber ball imaging (FBI, Jensen et al., 2016; Moss et al., 2019; Moss and Jensen, 2021) and fiber ball white matter modelling (FBWM, McKinnon et al., 2018). PyDesigner speeds up the tensor fitting regime by limiting computations within voxels that only contain brain tissue by using a brain mask. All aforementioned preprocessing steps are executed as part of the standard pipeline run.

Users can enable or disable image correction steps by parsing corresponding image correction flags instead of the --*standard* flag. Additional control flags are available to specify granularity of image corrections and tensor fitting. Information on all control flags is available at [PyDesigner - List of Flags](https://pydesigner.readthedocs.io/en/latest/reference/pydesigner_args.html). A completely pre-processed DWI using PyDesigner possesses minimal thermal noise and outliers and is co-registered to minimize motion. PyDesigner populates subject output directories with standard PyDesigner outputs.

### DESIGNER^1^

Preprocessed file from PyDesigner was first converted to MRtrix image format (*.mif*) using the function MRtrix3 function *mrconvert.* Then, all DKI-compatible *b*-value shells less than 3000 s/mm^2^ were extracted with *dwiextract* and parsed into DESIGNER’S tensor fitting function *tensorfitting.m* with the same tensor fitting parameters as PyDesigner (including IRLLS outlier detection and AKC correction) to generate standard DESIGNER output metrics. Fitting was performed with the default constraint *K_app_* > 0.

PyDesigner’s tensor fitting is adapted from DESIGNER by a straightforward Python translation of MATLAB code. Any differences seen in resulting maps are likely due to program dependent differences in the implementation of mathematical operations.

### DKE

Preprocessed images from PyDesigner (*dwi__preprocessed.nii*) were parsed through the MATLAB function *des2dke.m* (found in PyDesigner repository). This function extracts all *b*-value shells less than *b* = 3000 s/mm^2^, averages b0 volumes and concatenates them to form a DKE-compatible 4D NifTi file. This is done as DKE requires only a single b0 volume placed at the beginning of an input DWI for tensor fitting. This requirement limits accuracy of DKE’s linear least squares tensor fitting because a single data point is used to initialize the fit. DKE processed this file to generate standard diffusion and kurtosis parameter outputs. Additionally, DKE’s robustness in tensor estimation is limited as it does not perform any outlier detection or tensor correction. Fitting was performed with the default constraint *K_app_* > 0.

### DIPY

The same files used for DESIGNER processing were parsed into DIPY within a Python Jupyter Notebook. A DKI model was fitted to the data with *dipy.reconst.dki.DiffusionKurtosisModel()* using the default weighted least squares (WLS) and parameter values were extracted. DIPY is the only software among those tested that performs unconstrained tensor fitting.

### Postprocessing

A cerebral spinal fluid (CSF) excluded brain mask was created using FMRIB’s Automated Segmentation Tool (FAST) and *fslstats* with a brain-masked average b0 volume. This mask was applied to mean diffusivity (MD), fractional anisotropy (FA), and mean kurtosis (MK) maps to extract metrics values in non-CSF tissue. Voxels with MD ≤ 0, MK ≤ 0, and MK ≥ 10 were excluded as these are considered biologically implausible parameter values. These metrics were then compared across the four software to report on tensor fitting differences.

## RESULTS

All three commonly studied diffusion parameters (MD, FA, and MK) were found to be nearly identical in most voxels, especially for the FA and MD images, as seen in Figure 2 and 3. Differences between the software tools are more apparent with MK, particularly for the highly aligned fibers of the corpus callosum. Note that the PyDesigner MK appears to have more uniform intensity along the splenium of the corpus callosum, in comparison to the DESIGNER, DKE and DIPY estimates.

**Figure 2:**
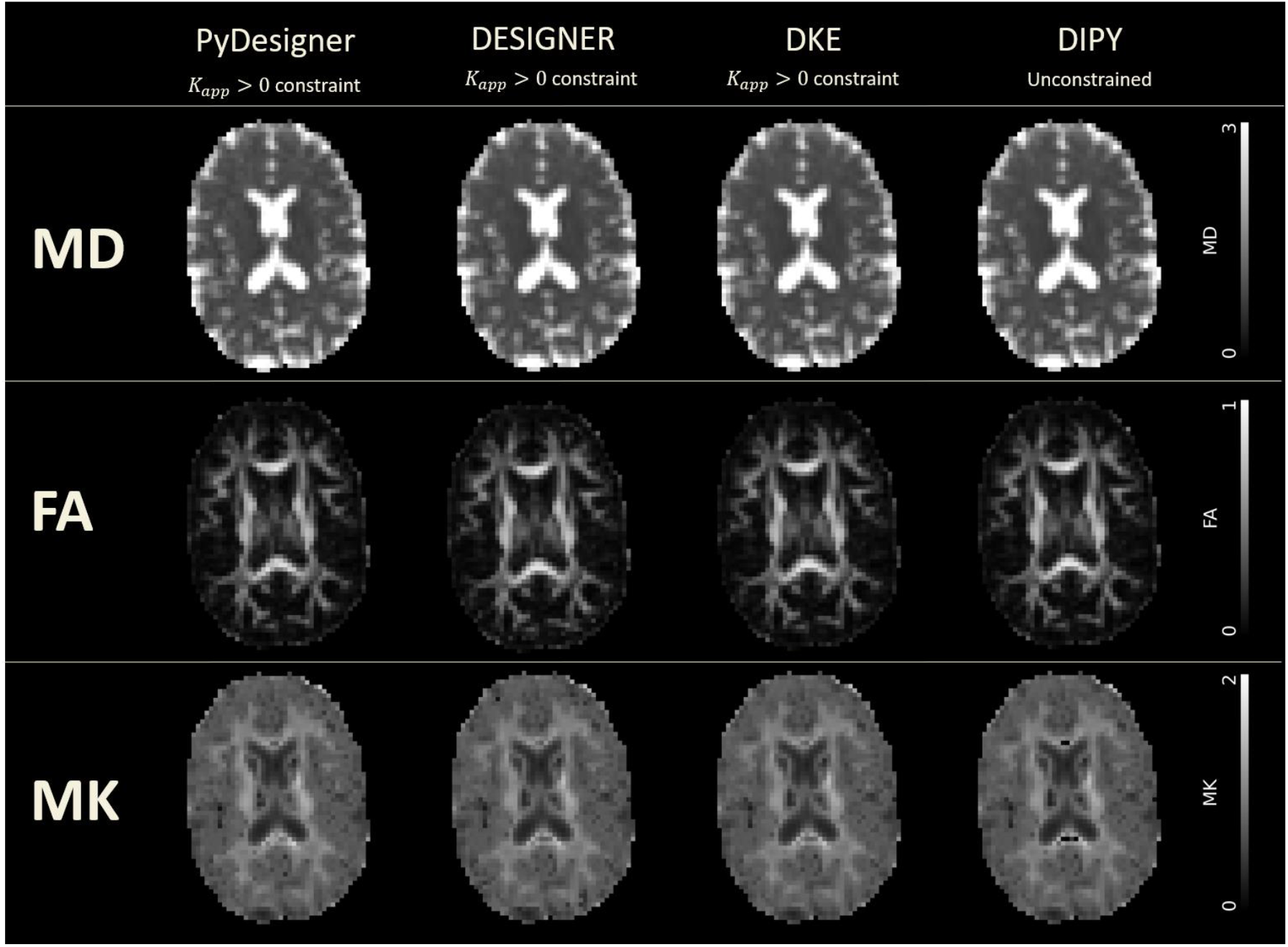
Commonly analyzed diffusion tensor and kurtosis imaging maps derived from PyDesigner, DESIGNER, DKE and DIPY. Tensor fitting was performed with *K_app_* > 0 constraint in PyDesigner, DESIGNER, and DKE, whereas unconstrained fitting was used in DIPY due to software limitations. The units for the MD scale bar are in μm^2^/ms, while the other scale bars are dimensionless.

**Figure 3:**
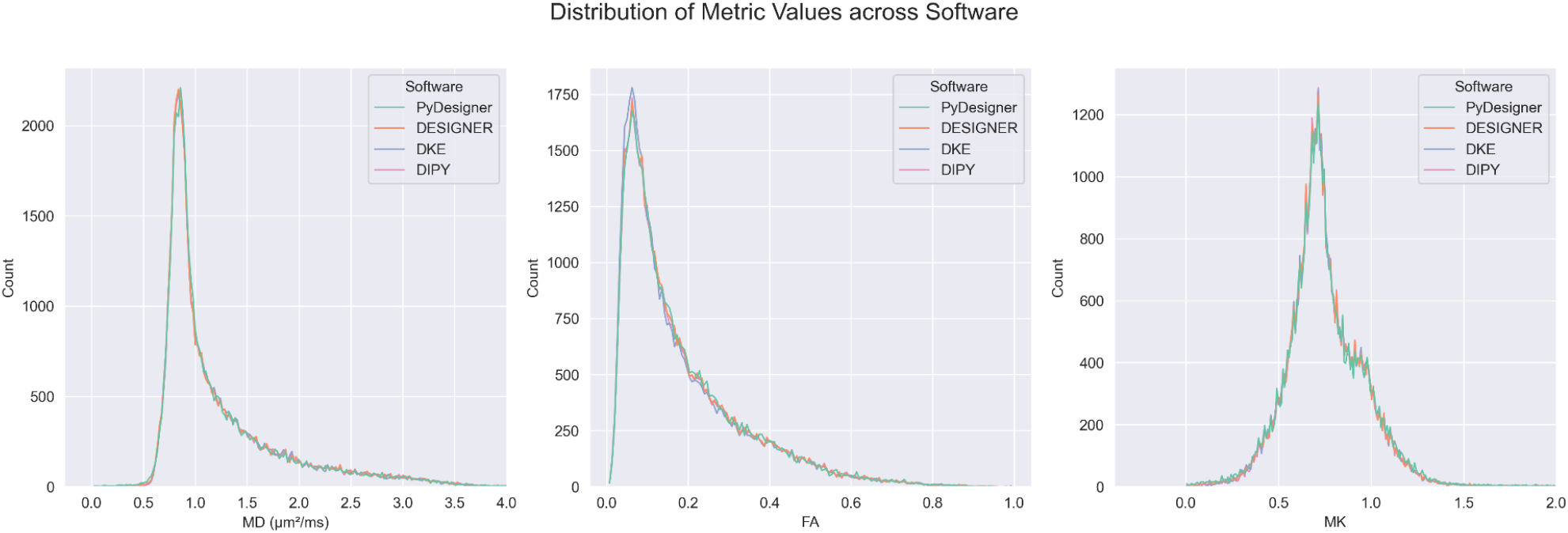
Distribution of computed values for FA, MD, and MK from PyDesigner, DESIGNER, DKE, and DIPY in cerebral spinal fluid (CSF)-excluded brain are similar across most voxels

Distribution plots of MD, FA, and MK, shown in Figure 3, display minimal differences between metrics across all four software. The MD, FA, and MK values are biologically plausible, except for a small number of voxels with MD exceeding the diffusivity of free water at 37 °C (3.0 μm^2^/ms), which likely reflects CSF partial volume effects. Inter-parametric correlations of MD vs. FA and MK vs. FA are shown in Figure 4 andFigure *5*, respectively. The MD and FA are nearly identical, with only minor discrepancies likely owing to differences in implementation of linear least squares fitting. For the MK vs. FA correlations of Figure 5, PyDesigner and DESIGNER again yield highly similar results, but deviations can be seen for DKE because, by default, it limits MK values to lie below 3 and with DIPY because it does not impose the *K_app_* > 0 constraint, resulting in more points with high FA together with low MK. Table 1 lists the Pearson correlation coefficients for the various comparisons. These again are quite similar, although the MD vs. FA correlation coefficient for DKE is somewhat larger than for the other three. Overall, the diffusion parameter estimates obtained from PyDesigner are consistent with those obtained with the other tensor fitting programs, thus providing supporting evidence of its accuracy and robustness.

**Figure 4:**
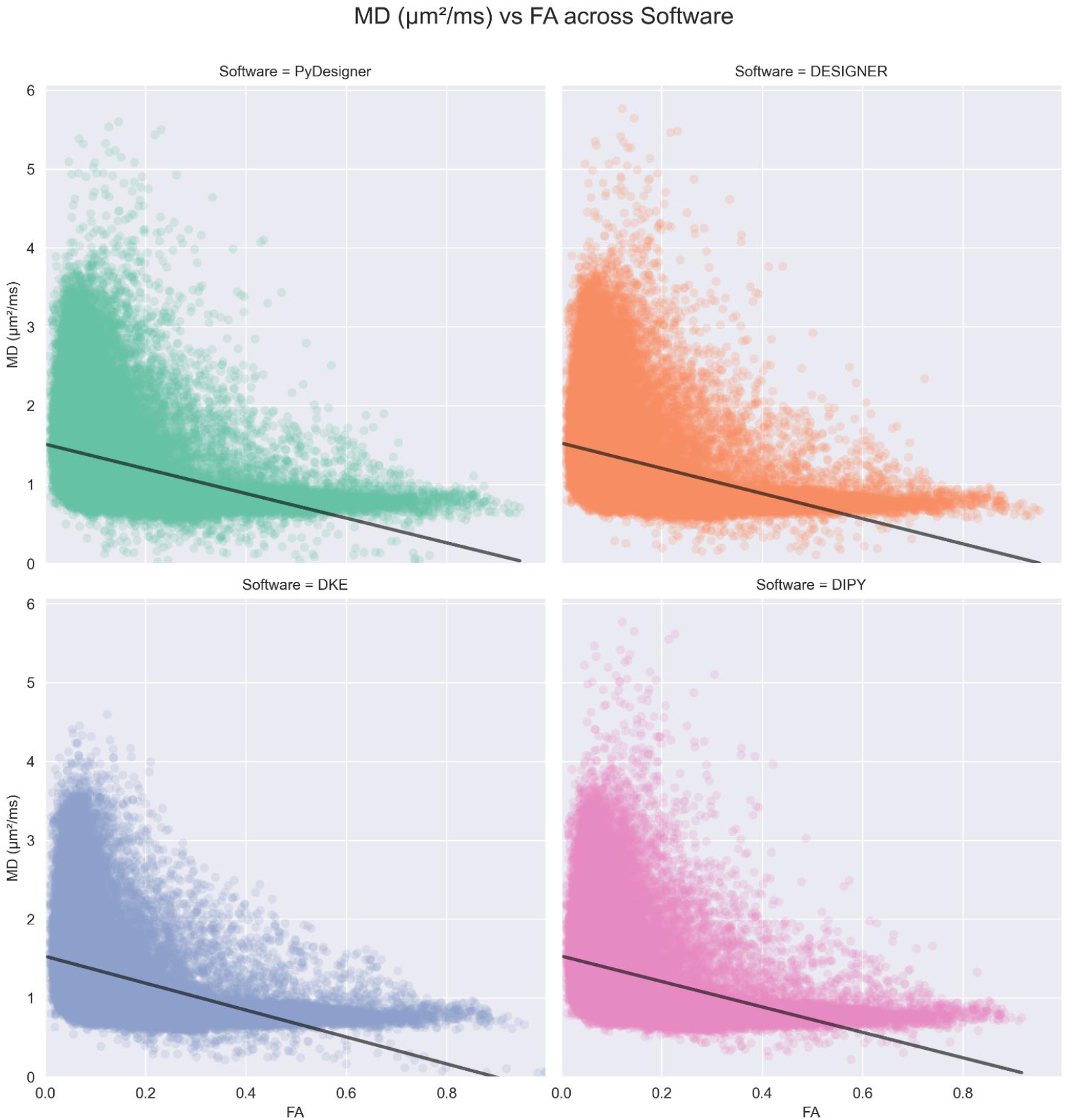
Plots of FA (x-axis) vs MD (y-axis) to illustrate the consistency of these diffusion parameters across the four software tools. Plots are sorted by software. The lines are best fits from linear regression.

**Figure 5:**
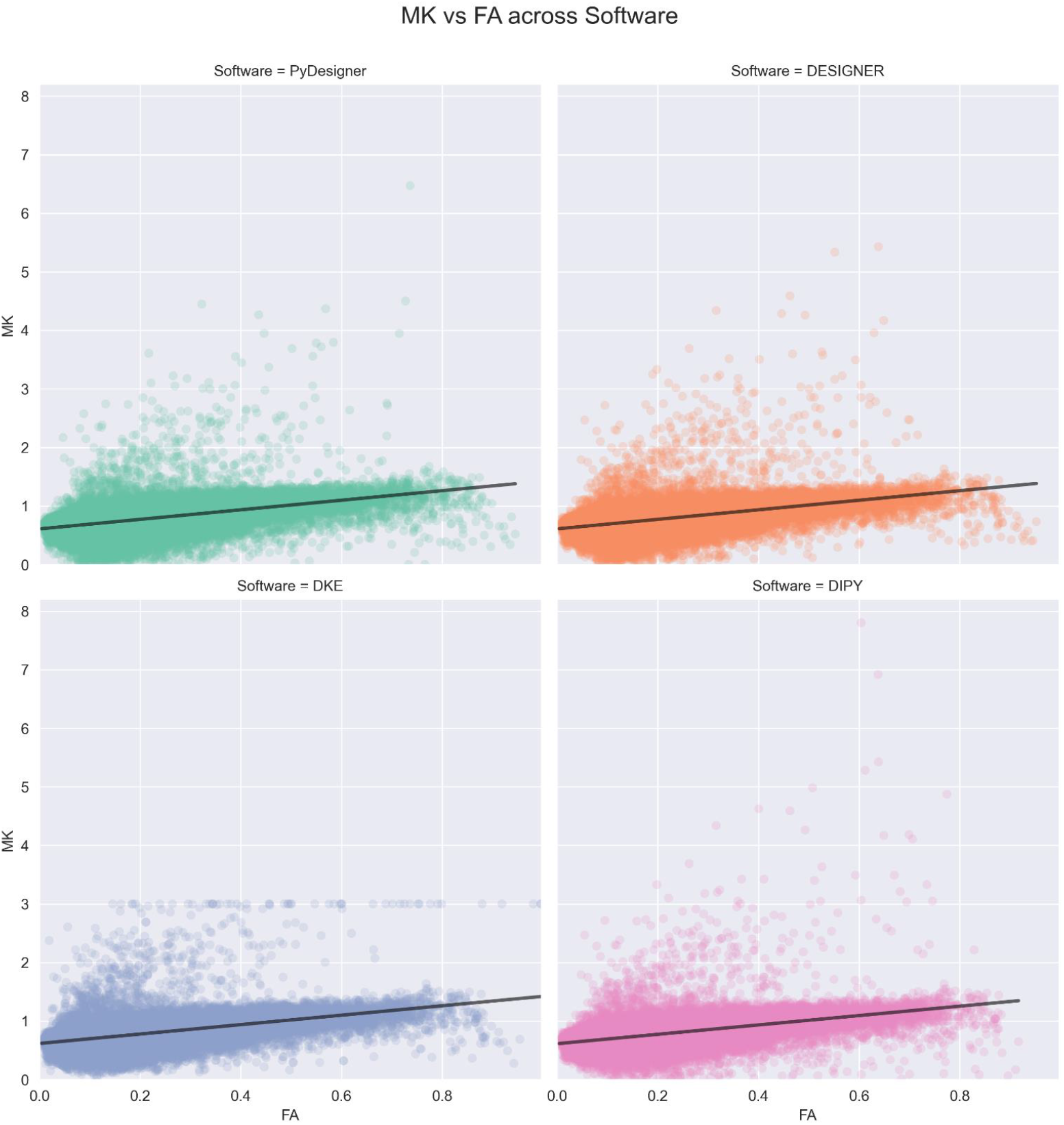
Plots of FA (x-axis) vs MK (y-axis) to illustrate consistency. Plots are sorted by software. The lines are best fits from linear regression. Note that the MK for the DKE calculations are restricted to be less than or equal to 3.

**Table 1:** Pearson correlation coefficients between FA and MD, and FA and MK across all four DKI analysis programs evaluated.

| **Software** | **Correlation** |  |
| --- | --- | --- |
|  | *FA with MD* | *FA with MK* |
| **PyDesigner** | -0.3786 | 0.5097 |
| **DESIGNER** | -0.3843 | 0.5153 |
| **DKE** | -0.4288 | 0.5156 |
| **DIPY** | -0.3854 | 0.5049 |

## DISCUSSION AND CONCLUSION

The primary motivation for developing PyDesigner was to implement the key elements of DESIGNER with all MATLAB code being replaced with Python, thereby allowing greater portability and accessibility. As our numerical results demonstrate, PyDesigner and DESIGNER yield nearly identical outputs. Nonetheless, there are a few additional options and default settings along with some minor bug fixes introduced while coding PyDesigner. These are described in detail in the online PyDesigner documentation (https://github.com/m-ama/PyDesigner). At the time of this writing, not all preprocessing features of DESIGNER such as B1 bias correction and DWI intensity normalization have been fully implemented in PyDesigner but are planned in future updates.

Here we also compared the PyDesigner tensor fitting calculations to those of the commonly used DKE and DIPY DKI analysis tools, showing that PyDesigner again yields similar results. Regarding the small differences that are found between PyDesigner and DESIGNER, on the one hand, relative to DKE and DIPY, on the other, we believe the two DESIGNER-based programs to be more accurate since they employ a more sophisticated fitting algorithm as discussed by Ades-Aron et al. (Ades-Aron et al., 2018). Combining constrained tensor fitting, outlier detection, and AKC correction yield robust and accurate tensor fitting seen in PyDesigner and DESIGNER.

A key advantage of PyDesigner over DESIGNER is that it is available in a Docker container called NeuroDock (dmri/neurodock), which greatly enhances portability and simplifies installation. This container runs across all major OS platforms compatible with Docker, including Microsoft Windows, Mac OS, and various Linux distributions. Docker’s container technology also enables straightforward deployment to high performance clusters (HPCs) for batch processing DWIs quickly on Docker-compatible local clusters, Amazon AWS, or Microsoft Azure.

PyDesigner also includes microstuctural modeling calculations that go beyond DKI, including White Matter Tract Integrity (WMTI) (Fieremans et al., 2011), FBI, and FBWM. For WMTI, a standard DKI dataset is adequate, and the associated microstructural parameters are calculated by default. However, it should be emphasized that the validity of WMTI is restricted to white matter regions with high FA (i.e., FA ≳ 0.4) and with some WMTI metrics having a limited accuracy due to assuming parallel alignment of axons in any given voxel. FBI (Jensen et al., 2016; Moss et al., 2019; Moss and Jensen, 2021) is a distinct dMRI method applicable throughout the cerebral white matter, which requires high *b*-value (i.e., *b* ≥ 4000 s/mm^2^) dMRI data sampled with a minimum of about 64 diffusion encoding directions (along with data for *b* = 0). The main outputs of FBI are the fiber orientation density function (fODF) for each white matter voxel, which can be used for white matter tractography and serves as an input for FBWM, as well as the intra-axonal fractional anisotropy (FAA). FBWM utilizes the dMRI data from both DKI and FBI to estimate the same parameters as WMTI but with improved accuracy. Thus, if this additional data is available, then FBWM estimates are preferred over those from WMTI (McKinnon et al., 2018). As with FBI, FBWM has only been validated in adult cerebral white matter.

Another notable feature of PyDesigner is multi-file input, which allows it to handle various file inputs - NifTi (.nii), compressed NifTi (.nii.gz), DICOM (.dcm), and MRtrix file format (.mif). PyDesigner is able to automatically identify acquisition information from header metadata regardless of input format and perform corrections accordingly, thereby supporting a hands-off approach. Regardless of differences in protocols, the same command (see above) can be used to process a wide variety of DWIs. PyDesigner thus saves time and effort by minimizing manual preprocessing steps and commands. In a recently released update (v1.0-RC10), this has been enhanced by introducing compatibility for multiple echo-time (multi-TE) datasets. This allows PyDesigner to run image preprocessing steps, which are largely independent of TE, on a multi-TE DWI to yield an image with minimal noise and artifacts. TE-dependent tensor calculations are then performed on each TE separately to produce diffusion or kurtosis metrics.

PyDesigner is still under development and improvements in existing features and the addition of new features are both expected in new updates. These will be detailed on the PyDesigner website (https://github.com/m-ama/PyDesigner), which provides both documentation and source code. Readers are encouraged to consult this website for the most up-to-date version of PyDesigner prior to beginning a new analysis. PyDesigner’s GitHub page also hosts a discussion forum where questions regarding PyDesigner can be submitted (https://github.com/m-ama/PyDesigner/discussions). The Docker implementation for portability is called NeuroDock (https://hub.docker.com/r/dmri/neurodock), which contains PyDesigner and its dependencies to enable processing across a wide array of platforms.

## FUNDING

Research reported in this publication was supported, in part, by National Institutes of Health grants R01AG054159, R01AG057602, R01AG055132, R01DC014021, R01NS110347, R21DA050085, F31NS108623, P20GM109040, P50DC000422, T32GM008716, and T32DC014435. Additional funding was provided by the Litwin Foundation.

## Footnotes

1 PyDesigner and DESIGNER rely on the same MRTrix3 and FSL tools and command syntax to perform image correction.

## Notes

### Competing Interest Statement

JHJ, JAH, and EF are co-inventors on US patents #8,811,706 and #9,965,962 that pertain to aspects of diffusional kurtosis imaging.

### Summary of Updates

Added funding information

